# Investigating metabolic interactions in a microbial co-culture through integrated modelling and experiments

**DOI:** 10.1101/532184

**Authors:** Aarthi Ravikrishnan, Lars M Blank, Smita Srivastava, Karthik Raman

## Abstract

Microbial co-cultures have been used in several biotechnological applications. Within these co-cultures, the micro-organisms tend to interact with each other and perform complex actions vis-à-vis a single organism. Investigating metabolic interactions in microbial co-cultures is crucial in designing microbial consortia tailored for specific applications. In this study, we present a pipeline integrating modelling and experimental approaches to understand metabolic interactions between organisms in a community. We define a new index named *Metabolic Support Index* (MSI), which quantifies the benefits derived by each organism in the presence of the other when grown as a co-culture. We computed MSI for several experimentally demonstrated co-culture systems and showed that MSI, as a metric, accurately identifies the organism that derives the maximum benefit. We also computed MSI for a commonly used yeast co-culture consisting of *Saccharomyces cerevisiae* and *Pichia stipitis* and observed that the latter derives higher benefit from the interaction. Further, we designed two-stage experiments to study mutual interactions and showed that *P. stipitis* indeed derives the maximum benefit from the interaction, as shown from our computational predictions. Also, using our previously developed computational tool MetQuest, we identified all the metabolic exchanges happening between these organisms by analysing the pathways spanning the two organisms. By analysing the HPLC profiles and studying the isotope labelling, we show that *P. stipitis* consumes the ethanol produced by *S. cerevisiae* when grown on glucose-rich medium under aerobic conditions, as also indicated by our *in silico* pathway analyses. Our approach represents an important step in understanding metabolic interactions in microbial communities through an integrating framework of modelling and experiments.

## 1. INTRODUCTION

Microbial co-cultures have been broadly used in several biotechnological applications, owing to their abilities to produce a broader pool of enzymes, which enable the synthesis or degradation of complex molecules. They also exhibit division of labour and provide a wider scope to leverage the joint metabolic capabilities of the organisms vis-à-vis a single organism. Microbial co-cultures have been employed to carry out complex functions ranging from xenobiotic degradation (1) to synthesising novel secondary metabolites (2, 3). More recently, the application of co-culture systems to produce biofuels has also been gaining traction, where groups of micro-organisms, either wild-type or engineered, have been employed to convert lignocellulosic biomass to ethanol (4–6).

Besides, there have also been advances in engineering microbial co-cultures to produce fine chemicals. Such synthetic microbial communities consist of organisms that have been engineered to communicate with each other via metabolic exchanges. For instance, in one such study (7), *Escherichia coli* cells were manipulated by engineering pathways to exchange metabolites for improving the titres of *n*-butanol. In a few other studies (8, 9), specialised metabolic pathways were engineered in separate *E. coli* cells to produce industrially important chemicals such as *cis, cis*-muconic acid, 3-aminobenzoic acid and resveratrol.

In addition to industrial applications, co-culture systems are also useful to understand the interactions happening between the organisms. The organisms as a community, orchestrate several complex functions by communicating with one another, commonly via metabolic interactions (10). These interactions define the relationships between the organisms and shape the overall structure of the microbial community. Additionally, computational studies have also provided insights into the pairwise relationships between the organisms in a community (11–16).

Understanding the metabolism of individual organisms, as well as investigating the metabolic interactions that happen in a community is central to *designing* a microbial consortium for a given application. There are likely multiple metabolites being exchanged between the micro-organisms in a consortium. Experimentally identifying and understanding the role of these metabolic exchanges and interactions entails the profiling of both intracellular and extracellular metabolites. Such untargeted metabolomic profiling becomes very difficult (17), especially when it involves a co-culture of organisms, owing to the characterisation of organisms in the co-culture, although complete exometabolomic analysis has been recently carried out for a few microbial communities (18, 19).

In this study, we design a comprehensive workflow integrating computational and experimental approaches to predict and validate the microbial interactions in a co-culture. We adopt a three-pronged approach to study a co-culture of two industrially important yeast species, viz. *Saccharomyces cerevisiae* and *Pichia stipitis*, drawing on computational analyses, physiological studies and ^13^C-labelling experiments. First, using our previously developed graph-theoretic algorithm, MetQuest (20), we quantify the possible benefits derived by the micro-organisms when they stay together in a community. We propose a new metric called “Metabolic Support Index (MSI)” to determine which organism derives a relatively higher benefit from the interaction. We show that this metric can successfully quantify the interactions between the organisms using a few examples of the previously demonstrated microbial co-cultures. We then identify the *metabolic exchanges* happening between *S. cerevisiae* and *P. stipitis* and experimentally demonstrate the predicted metabolic interaction by performing experimental verifications. To this end, we perform physiological experiments to identify the organisms that benefit this interaction. Lastly, we perform isotope-labelling studies to determine the metabolic interactions between the organisms and understand the pathways where the exchange metabolite is involved. The results from our workflow on this model co-culture system indicate that *P. stipitis* benefits from the interaction through the uptake of ethanol produced by *S. cerevisiae* when grown on glucose-rich medium under aerobic conditions. Our approach represents an important step in integrating modelling with experiments to understand and characterise microbial interactions in communities and underlines the utility of metabolic modelling to understand, and possibly design microbial communities for specific applications.

## 2. RESULTS

In this study, we present a pipeline integrating modelling and experimental approaches to understand the microbial interactions between organisms in a community. As the first step of this pipeline, we use our previously developed graph-theoretical approach, MetQuest (20) to study the community metabolic network, and compute the *Metabolic Support Index* (MSI). This index quantifies the benefits derived by each organism in the presence of the other when grown together as a co-culture. We calculate MSI for several experimentally demonstrated co-culture systems and show that MSI, as a metric, accurately identifies the organism that derives the maximum benefit. Further, we design two-stage experiments to study mutual interactions and show that *P. stipitis* indeed derives the maximum benefit from the interaction, as shown from our computational predictions Also, we identify the metabolic exchanges happening between *S. cerevisiae* and *P. stipitis* by analysing the pathways spanning the two organisms. Further, using the HPLC profiles and isotope labelling studies, we show that *P. stipitis* consumes the ethanol produced by *S. cerevisiae* when grown on glucose-rich medium under aerobic conditions, as also indicated by our *in silico* pathway analyses.

### 2.1 *Metabolic support index* quantifies the benefits derived by organisms in a community

To quantify the benefits derived by the organisms in a community, we compute a metric, *Metabolic Support Index* (MSI), as described in Methods. MSI essentially quantifies the fraction of an organism’s metabolic network that is ‘enabled’ by the presence of the other organism. An MSI of zero indicates that the organism does not receive any metabolic support from the other, while an MSI of unity indicates that the organism receives all the metabolites required to ‘enable’ the stuck reactions. We calculated MSI for several pairs of organisms, which have been experimentally demonstrated to grow together as a community and where mutual interactions between the organisms have been reported (Table 1). By comparing the MSI of individual organisms, we determine the organism from the pair that derives the maximum benefit. In addition, we also quantify the interactions based on the increase in the number of metabolites produced by one organism in the presence of the other (Supplementary Table 1). Such an increase points to the synergy and the possible metabolic interactions between the organisms. These observations are also in agreement with those reported previously (11).

**Table 1:** Metabolic support index to quantify pairwise interactions –. This table shows the MSI of two organisms calculated for different pairs of organisms that have been experimentally studied. The two organisms in a co-culture are marked Organism A and Organism B for ease of representation. Higher the value of MSI, better is the benefit the organism derives through this interaction. The references in Column 2 and 3 represent the source of the genome-scale metabolic model. The references in Column 6 represent the experimental studies where the co-cultures of organisms have been studied. We observe that amongst every pair of organisms, the organisms in Column 2 derive higher benefit (shown in boldface font), in agreement with previously reported experiments (Column 6).

| S. No | Organism A | Organism B | MSI (A\|AUB) | MSI (B\|AUB) | Comments |
| --- | --- | --- | --- | --- | --- |
| 1 | *Ketogulonicigenium vulgare* (31) | *Bacillus megaterium* (31) | **0.107** | 0.0 | *B. megaterium* is a helper strain for *K. vulgare* (38, 39) |
| 2 | *Clostridium cellulolyticum*(32) | *Clostridium acetobutylicum* (34) | **0.039** | 0.001 | *C. acetobutylicum* helps *C. cellulolyticum* when grown together in a cellulose-rich medium (40) |
| 3 | *Pichia stipitis* (35) | *Saccharomyces cerevisiae* (41) | **0.063** | 0.025 | *P. stipitis* benefits the interaction with *S. cerevisiae* (this study) |
| 4 | *Yarrowia lipolytica* (36) | *Cellulomonas fimi* (36) | **0.119** | 0.014 | *C. fimi* provides additional metabolites to *Y. lipolytica* in a co-culture setup (18) |
| 5 | *Desulfovibrio vulgaris* (36) | *Methanococcus maripaludis* (36) | **0.176** | 0.03 | *D. vulgaris* benefits the interaction with *M. maripaludis* (42) |

In all the cases considered, organism A (Column 2) derives a higher benefit in the co-culture than organism B (Column 3). These observations are in exact agreement with those made by the experimental studies (Column 6), evaluating the relative biomasses of the two organisms and their ability to co-exist. For instance, we observe that the MSI of *Ketogulonicigenium vulgare* is 0.1, while that of *Bacillus megaterium* is 0. This indicates that no additional pathways have been activated in *B. megaterium*, in the presence of *K. vulgare.* On the other hand, in *K. vulgare*, 41 additional reactions have been activated in the co-culture (Supplementary Table 1), which could otherwise not take place in the monoculture. These results indicate the additional metabolic support *K. vulgare* receives from *B. megaterium*. It is interesting to note that this co-culture system has been widely studied for its applications in Vitamin C production; these studies also indicate that *B. megaterium* is a helper strain that enhances the growth and proliferation of *K. vulgare* (21–23).

In another co-culture consisting of *Yarrowia lipolytica and Cellulomonas fimi*, we observe that MSI of *Y. lipolytica* (0.119) is almost 10 times that of *C. fimi* (0.014). This is because of the additional 123 reactions that have been activated in *Y. lipolytica* in the presence of *C. fimi.* Activation of these reactions also points to the enrichment in the metabolic capabilities of the *Y. lipolytica*, as also observed from the increase in the *scope* size of joint metabolic networks (Supplementary Table 1). These results indicate that *Y. lipolytica* derives the maximum benefit when grown as a co-culture with *C. fimi.* In sum, these results indicate the overall *metabolic* benefit that *Y. lipolytica* receives, which could have led to the increase in its growth in co-culture, as observed in the experimental studies previously carried out (18). Similarly, in all the other cases we have considered, we show that the trends in MSI agree exactly with the results demonstrated experimentally (Table 1).

Next, we calculated the MSI for the co-culture system of our interest consisting of *S. cerevisiae* and *P. stipitis.* Here, we observe that MSI of *P. stipitis* is 0.063, nearly three times more than that of *S. cerevisiae* (0.025). In addition, we observe 21 reactions in *P. stipitis* that have been enabled in the presence of *S. cerevisiae.* These results indicate that *P. stipitis* benefits the interaction with *S. cerevisiae*, which we proceeded to verify using growth kinetic experiments.

### 2.2 In silico pathway analyses reveals several interesting metabolic exchanges

To investigate the metabolic exchanges between *S. cerevisiae* and *P. stipitis*, we exhaustively enumerated all the pathways on a *community metabolic network*, using our previously developed algorithm MetQuest (see Methods). MetQuest identifies all possible pathways from a given set of seed metabolites to all the *reachable metabolites*, whose size is less than or equal to the cut-off. The pathway so obtained, is complete, i.e., it contains all the reactions necessary to produce every metabolite constituting the pathway.

We computed all the pathways starting from D-Glucose in *S. cerevisiae* that lead to various metabolites in *P. stipitis*. We analysed these pathways for the presence of metabolites from *S. cerevisiae* that resulted in the production of target metabolites in *P. stipitis.* In total, we observed pathways for 668 metabolites produced by *P. stipitis.* Of these, pathways to 634 metabolites consist of exchange metabolites from *S. cerevisiae* (Supplementary Table 2). Further, from all these pathways, we observed a total of 45 metabolites from *S. cerevisiae*, which are involved in the pathways producing various metabolites in *P. stipitis*. Of these 45 metabolites, we note that acetaldehyde, α-ketoglutarate, ethanol and sorbitol were the most commonly exchanged metabolites, which were involved in the production of over 400 metabolites in *P. stipitis* (Supplementary Figure S1).

Further, to determine if there were any additional benefits in terms of metabolic exchanges leading to the overall biomass production, we analysed all the pathways from *S. cerevisiae* D-glucose to the amino acids in *P. stipitis.* Amongst the 45 metabolites listed above, we observed a total of 34 exchange metabolites that formed a part of the pathways producing different amino acids (Supplementary Table 3). It was interesting to note that of the most commonly exchanged metabolites, acetaldehyde, alpha-ketoglutarate, ethanol and sorbitol take part in the production of over ten amino acids.

Several studies previously carried out on this coculture have demonstrated its ability to produce ethanol from a variety of substrates (24, 25). However, from our computational predictions, we observed ethanol transport from *S. cerevisiae* and *P. stipitis*, which participated in the production pathways of 418 metabolites. We then decided to experimentally check if ethanol was indeed exchanged between the organisms, as predicted from our pathway analyses. This would provide us with insights on designing better processes where a co-culture of these organisms is used for ethanol production.

### 2.3 *P. stipitis* exhibits a higher growth in the cell-free supernatant of *S. cerevisiae*

To check the existence of mutual interactions between *S. cerevisiae* and *P. stipitis*, we carried out mono-culture and co-culture growth kinetics experiments with these organisms on a minimal medium containing 10 g/L D-glucose as the carbon source (for both the sets of experiments). From the growth curves (Figure 3), we observed that the co-culture cell density was in between that of the respective monocultures. This provided indications on the ability of the organisms to co-exist. Further, the co-culture showed a diauxic pattern of growth curve, indicating that a few metabolites from the supernatant may serve as the carbon source.

**Figure 1:**
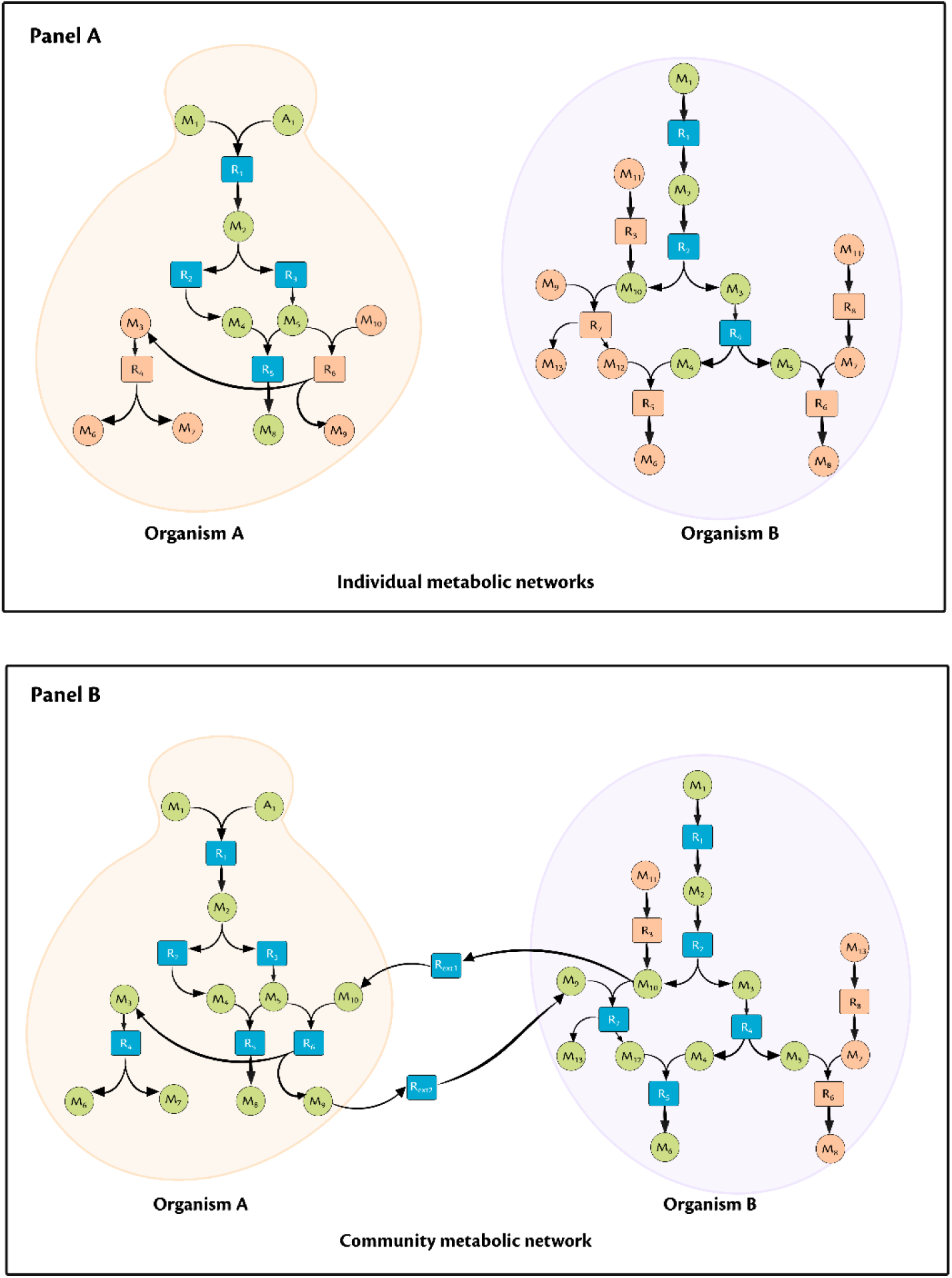
Metabolic Support Index (MSI) calculations. *Panel A - Bipartite graph of individual organisms, Panel B - Community metabolic network showing the metabolic exchanges.* Circular nodes are metabolites, square nodes are the reactions, green colour circular nodes represent the metabolites in the scope of M1 and A1, orange colour circular nodes represent the metabolites that is not present in the scope, orange colour square nodes represent the stuck reactions, blue colour square nodes represent the visited nodes. Here we see that *n*_*stuck,A*|*A*_ = 2, *n*_*stuck,A*|*A*∪*B*_ = 0, *MSI*(*A*|*A* ∪ *B*) = 1; similarly *n*_*stuck,B*|*B*_ = 5, *n*_*stuck,B*|*A*∪*B*_= 3, *MSI*(*B*|*A* ∪ *B*) = 0.4. Thus, we see that the Organism A benefits from this interaction with Organism B.

**Figure 2:**
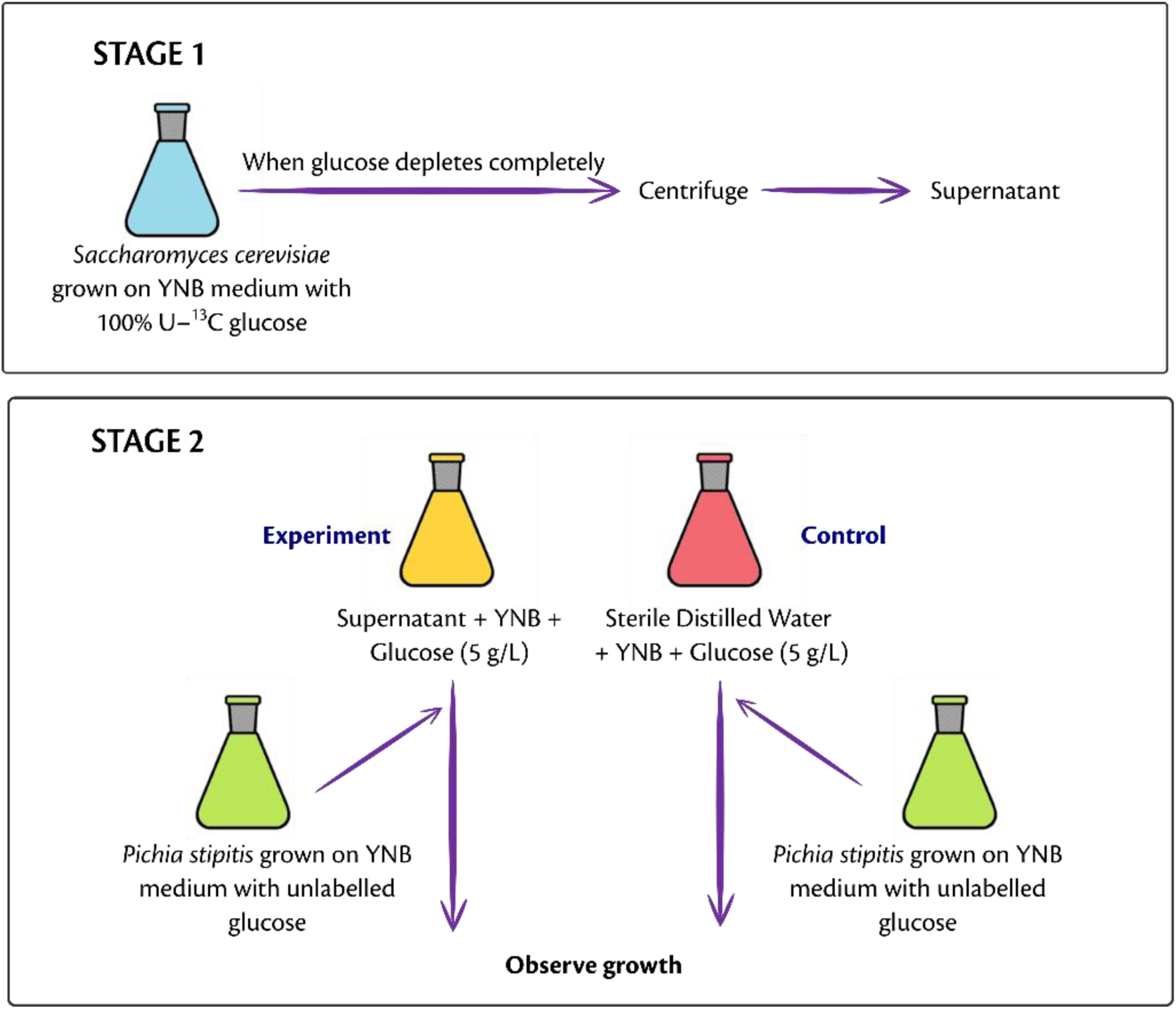
Two-stage experimental setup to identify metabolic interactions. In stage 1 (top panel), the first organism is grown in a minimal medium consisting of glucose. In stage 2 (bottom panel), the supernatant from the first organism is used as the medium to grow the second organism and the growth is compared with that of the control where sterile distilled water is used instead of the supernatant.

**Figure 3:**
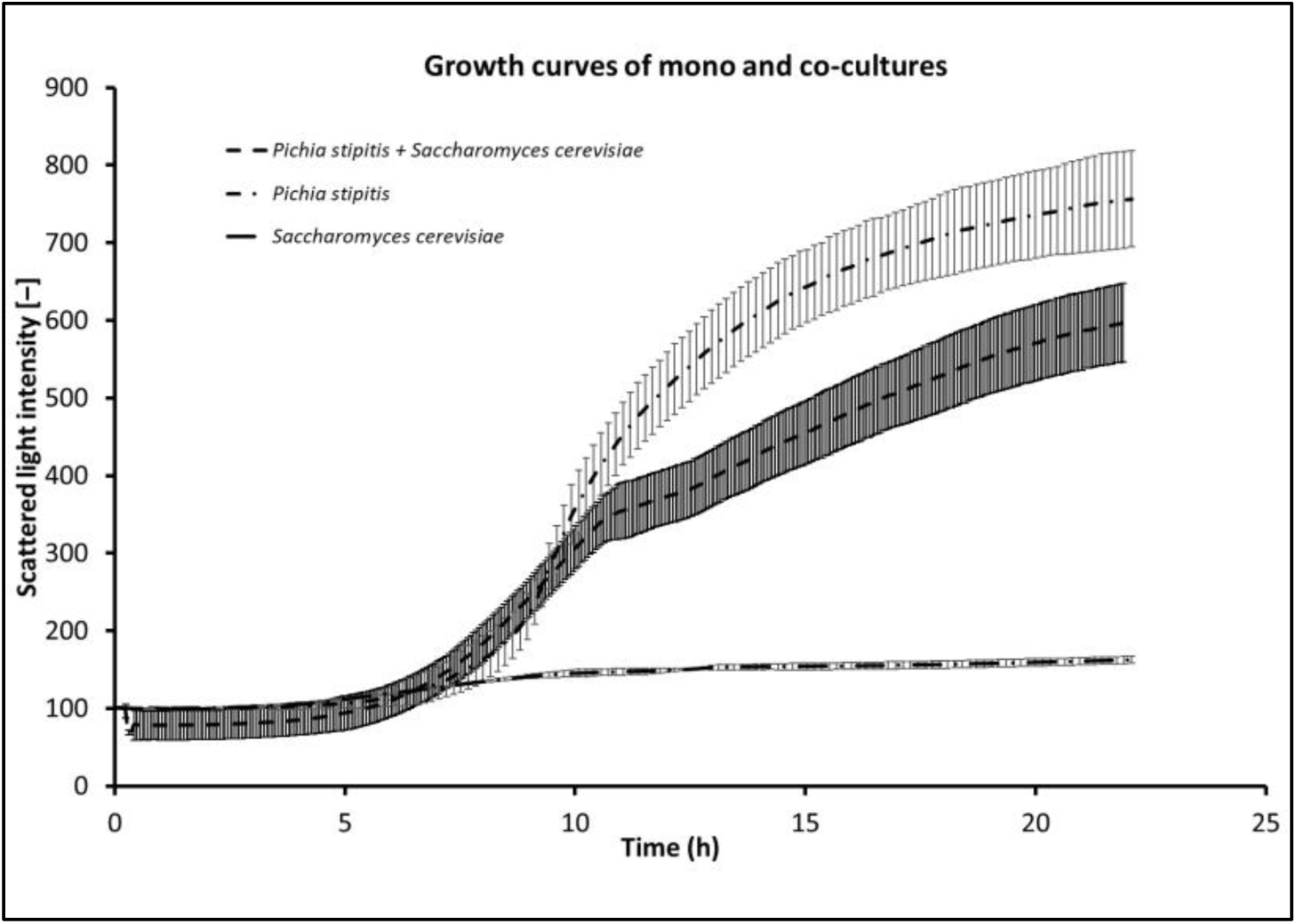
Growth curves of mono-and co-cultures of *Pichia stipitis* and *Saccharomyces cerevisiae*. Note that the growth curve of co-culture exhibits a diauxic pattern and lies between that of mono-cultures of *P. stipitis* and *S. cerevisiae*.

Next, to identify the organism that benefits the interactions, we designed and carried out the two-stage experiment, as described in methods. Interestingly, from the growth curves of second stage (Figure 4), we observed that at the end of 24 h, *P. stipitis* exhibited a 1.34-fold higher cell density (518.15) when grown in the supernatant of *S. cerevisiae*, in comparison to that seen in the control (309.41) at the end of 24 h. In addition to the improved cell density, *P. stipitis* clearly exhibited a diauxic growth pattern, indicating the presence of an alternate metabolite in the supernatant that was used as the carbon source. However, we did not observe any increase in the cell density of *S. cerevisiae* when grown in the supernatant of *P. stipitis.* This observation also corroborates the higher MSI for *P. stipitis* in the community, indicating that *P. stipitis* derives a higher benefit when grown together with *S. cerevisiae* under these specific conditions.

**Figure 4:**
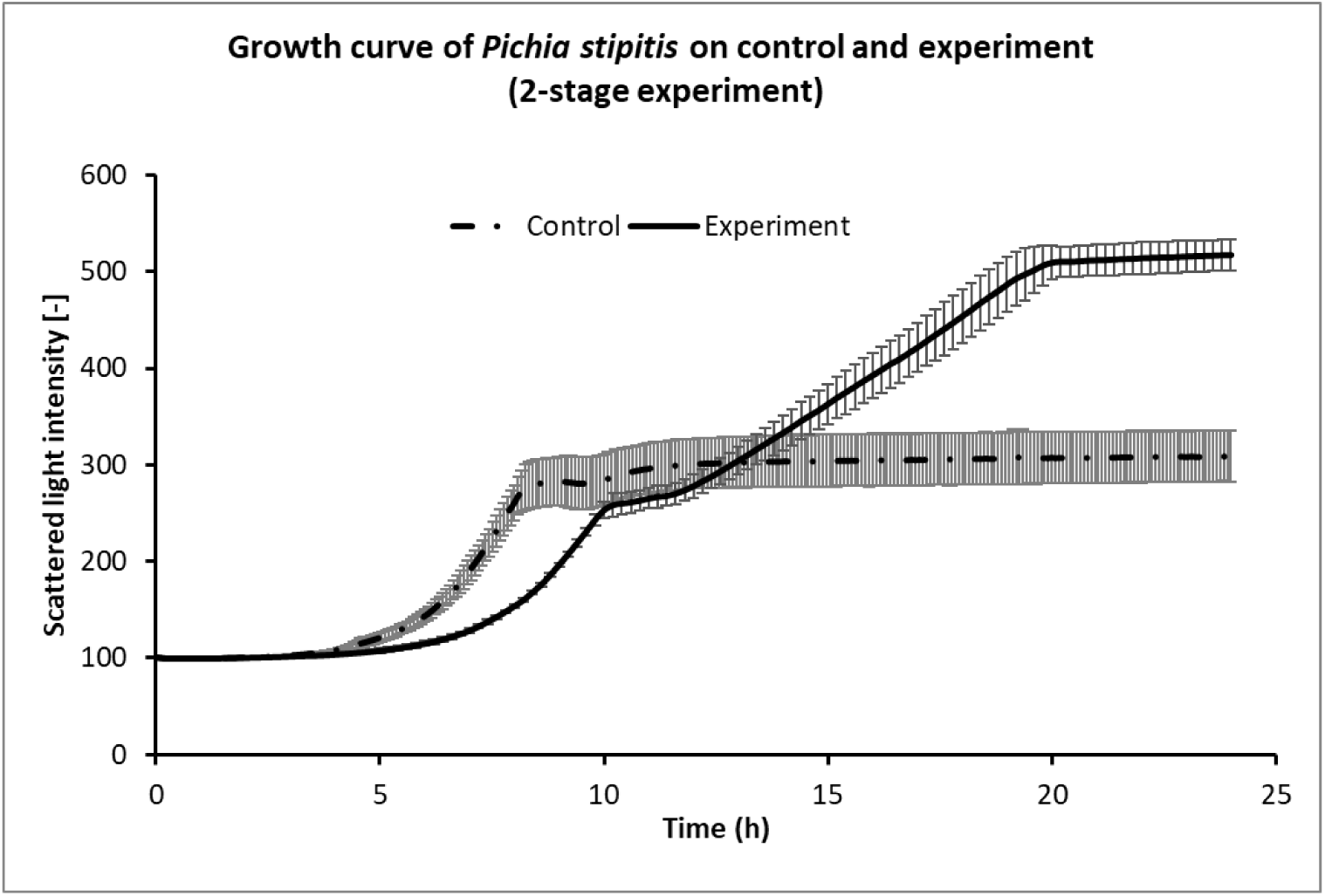
Comparison of the growth curve of *P. stipitis* grown on the supernatant of *S*. *cerevisiae* with that grown in control. The graph shows the consensus curve for the triplicates of control and experiment. Note the diauxic pattern of growth curve when *P. stipitis* is grown in the supernatant of *S. cerevisiae*.

### 2.4 *In silico* pathway analyses and Isotope labelling studies confirm ethanol transport from *S. cerevisiae* to *P. stipitis*

In the next step, we experimentally determined the metabolic exchanges happening between *S. cerevisiae* and *P. stipitis* using the same two-stage experimental setup. We performed HPLC analyses on the samples withdrawn at 0 h and 24 h timepoints of the second stage, where *P. stipitis* was grown in the supernatant of *S. cerevisiae*. Interestingly, we observed that the concentration of ethanol had reduced from 2.44 g/L (in 0 h) to 0 g/L (24 h).

Motivated by the reduction in the ethanol concentration and the predictions of metabolic exchanges from our computational studies (Sec 2.2), we hypothesised that *P. stipitis* was consuming the ethanol produced by *S. cerevisiae.* To confirm this, we studied the ^13^C-labelling patterns of the amino acids arising from ethanol metabolism. We used the metabolic network of *S. cerevisiae* to understand the ethanol metabolism in *P. stipitis* (26). From this, we identified the sets of amino acids arising directly and indirectly from ethanol metabolism (Supplementary Figure S2). The former set includes aspartate, valine and alanine. The latter set includes serine, derived from 3-phosphoglycerate that arises from gluconeogenesis, which is one of the indispensable pathways when yeasts grow on nonfermentable carbon sources (27). We then checked for the incorporation of ^13^C-labels in these amino acids specifically in the (M-57)+ fragment of the respective amino acids. Interestingly, in all these amino acids, we observed isotopomers with a higher fraction of ^13^C incorporated (Figure 5). For instance, from the pool of amino acid aspartate, we find 28% of m4 isotopomer where all the four carbon atoms carry a ^13^C label. In addition, we also noted that there is a negligible fraction of m1 isotopomer in all the four amino acids. Further, we observed that the average carbon labelling (0.39) in these amino acids is significantly higher than that found in the control (0.015), indicating the uptake of a labelled metabolite from the supernatant (Supplementary Table 4). In addition, the average carbon labelling observed in the amino acids (0.39) was almost equal to the fraction of labelled substrate present in the supernatant (0.42 ± 0.01) indicating that nearly all the observed labels in the amino acids originated from ethanol.

**Figure 5:**
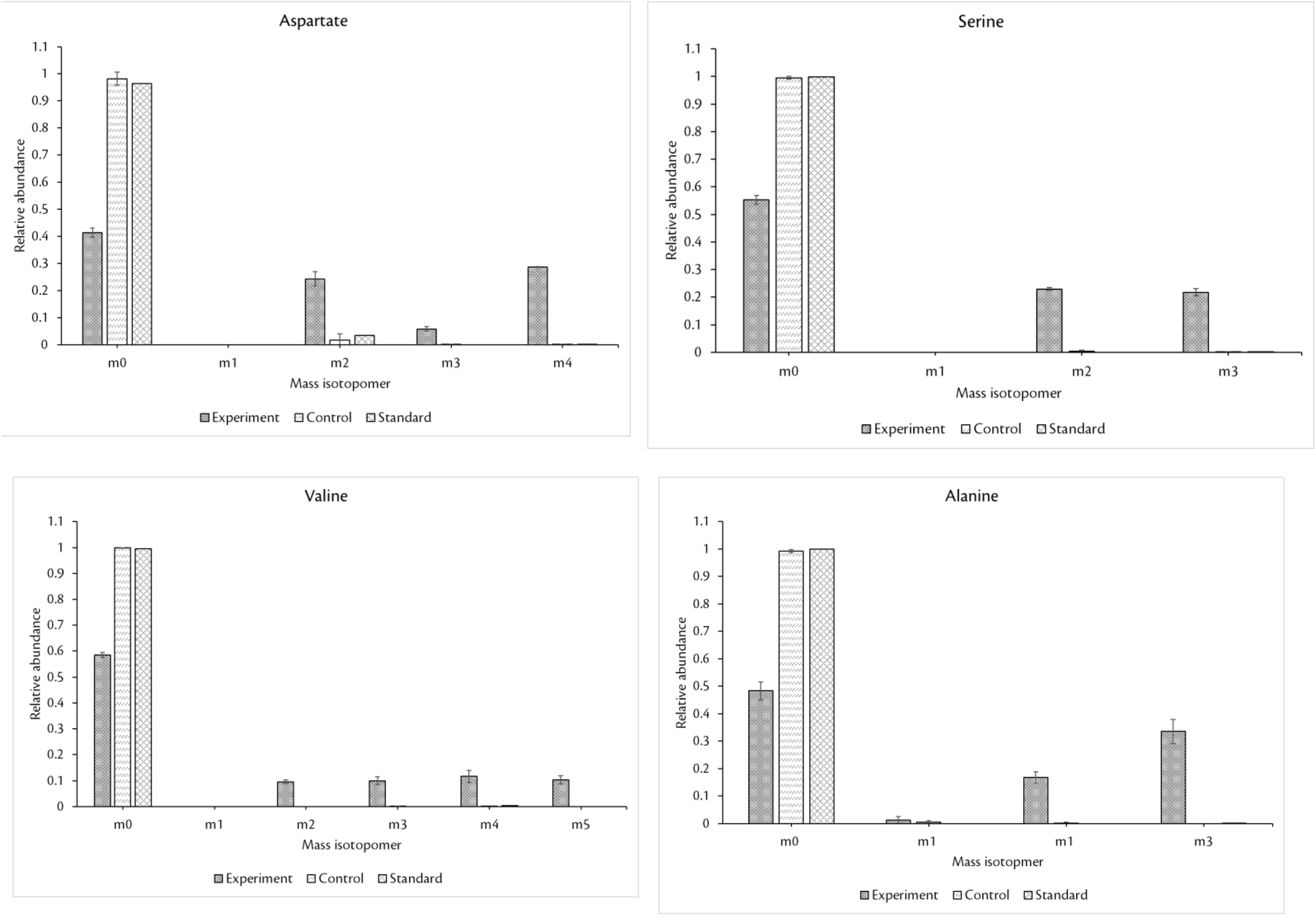
Mass isotopomer distributions of four different amino acids arising from ethanol metabolism. The relative abundance of heavier isotopomers of amino acids is much higher in comparison to that of control, indicating the uptake of a labelled carbon source. Note that we considered (M-57)+ fragment for our analysis. The three bars indicate the results from experiment, control and the standard, as indicated in the legend.

## 3. DISCUSSION

Microbial co-culture systems have been used for several biotechnological applications, where the metabolic capabilities of two different organisms have been exploited. Several recent studies have designed co-culture systems with micro-organisms that have been genetically modified to exchange metabolites with one another. Moreover, these microbial co-cultures also serve as a model system to study the fundamental cell-cell communication. There has been a growing interest in studying these microbial co-cultures/communities and identifying the interactions therein.

In this study, we systematically understand the metabolic interactions between the micro-organisms in a co-culture by integrating modelling and experimental approaches. To this end, we applied our previously developed graph-theoretic algorithm that operates on the metabolic networks of the micro-organisms. We defined a new metric termed *Metabolic Support Index* (MSI) that quantifies the support each organism receives from the other, in a community. We computed this metric on several co-culture systems that have been experimentally studied so far, demonstrating that this metric can correctly identify the beneficiaries in the microbial interactions.

A few other studies incorporate the nutrient conditions and quantify the cooperation between the organisms in a community. These studies identify the synthesising capabilities of “joint metabolic networks” (11, 28) and identify the benefits in terms of the metabolites produced by the individual organisms in a community. Our metric, MSI, on the other hand, determines the number of additional reactions that can be activated in one organism due to the presence of other, in addition to the metabolic enrichment. Furthermore, analyzing these reactions and identifying the transporters can provide insights on targets for over-expression. These targets can be used to improve the cooperativity between the organisms.

Further, to integrate the predictions with the experiments, we calculated MSI for a well-known yeast co-culture system consisting of *P. stipitis* and *S. cerevisiae* and identified that *P. stipitis*, with a higher MSI, benefits in this interaction. Moreover, we also identified the pathways spanning the two organisms and identified the set of metabolites that can be exchanged between these organisms. We also performed two-stage experiments using this co-culture and found that: (a) *P. stipitis* benefits the interaction, as shown by a higher value of MSI compared to that of *S. cerevisiae* and (b) *P. stipitis* consumes the ethanol produced by *S. cerevisiae.* Also, from the labelling studies, it is also interesting to note the complete absence of m1 isotopomer in all the amino acids analysed.

Our deconvoluted two-stage experiment is a simple methodology to establish a *proof-of-concept* for identifying the microbial interactions in a community. Indeed, the dynamics of the organisms in the co-culture may be different from that observed when the organisms are independently cultivated on the cell-free supernatants. However, deciphering the co-culture dynamics would entail the use of advanced molecular biological techniques. Moreover, isotope labelling patterns of the individual organisms from the co-culture experiments are difficult to obtain. Although recent studies seek to identify labelling patterns in co-culture, these methods either require extraction of a large number of peptides (29) or are restricted to a co-culture of organisms with identical biomass composition (30).

In addition, our computational method to determine the relationship between the organisms in a co-culture, captures only the positive interactions between the organisms, as with other graph-based method based on network expansion (11). Weighting the pathways based on their importance could help in further improving the power of MSI to predict interactions.

In sum, our study provides a systematic methodology to understand the interactions in a microbial community by integrating computational and experimental paradigms. We believe that such a model-integrated approach would provide us a better understanding of the intricate metabolic interactions happening between the organisms in a community. Our study also underlines the utility of computational analyses to generate testable hypotheses regarding the interactions between various microbes. This generic framework can be extrapolated to study the metabolic interactions in many microbial communities. Overall, our integrated approach can pave the way for rationally designing and engineering microbial consortia tailored towards specific industrial applications.

## 4. MATERIALS AND METHODS

### 4.1 Computational Analyses

#### 4.1.1 In silico calculation of MSI to identify the benefits derived by the organisms

To quantify the benefits derived by the organisms in a community, we define a metric termed *Metabolic support index* (MSI). For this, we used our previously developed algorithm MetQuest (20) to determine the number of reactions that can be *visited* or *stuck*, depending on the presence of precursor metabolites. As defined previously, *stuck* reactions are those reactions whose precursor metabolites cannot be synthesised by the metabolic network using the given input conditions, while *visited* reactions indicate those reactions whose input metabolites are present and hence the reaction can proceed. We define MSI as the fraction of blocked/stuck reactions relieved in organism A by the presence of another organism B (Figure 1), i.e.,

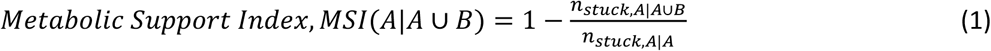

where *n*_*stuck,A*_ denotes the number of reactions that are stuck in the (bi-partite) metabolic network corresponding to organism A,

*n*_*stuck,A*∪*B*_ denotes the number of reactions that are stuck in the bi-partite metabolic network corresponding to the community metabolic network of A and B, and

*A* ∪ *B* denotes the community metabolic network comprising both organisms A and B

*MSI*(*A*|*A* ∪ *B*) = 1 indicates that organism A fully benefits in the interaction with organism B

*MSI*(*A*|*A* ∪ *B*) = 0 indicates that organism A does not derive any benefit from the interaction with organism B

To calculate MSI for a micro-organism, we first constructed the directed bipartite graph of the individual organism, followed by the joint bipartite graphs of combination of organisms from their respective genome-scale metabolic models using the *construct*_*graph* module in MetQuest python package. Depending on the availability of the genome-scale models of the organisms, we obtained them either from the respective publications (31–35) or from the Path2Models database (36). We applied our algorithm using a set of starting *seed* metabolites, which included the cofactors, coenzymes and the carbon source used in the respective experiments (see Supplementary dataset). For our co-culture of interest, i.e., *S. cerevisiae* and *P. stipitis*, the seed metabolite set consisted of components from YNB minimal medium, D-glucose as the carbon source along with the set of cofactors and coenzymes.

Next, using the *guided*_*bfs* module in the MetQuest package, we obtained the number of “stuck” reactions from two different scenarios: (a) when the organisms are analysed independently as “single” graphs (*n*_*stuck,A*|*A*_), (b) when they are grown along with other organisms as a “joint” graph (*n*_*stuck,A*|*A*∪*B*_). Using the equation (1), we calculate the MSI of the organisms. In addition, in both the cases, we also obtained the *scope*, i.e., the set of metabolites that can be produced from the given set of seed metabolites. All the simulations were carried out in Python 3.6 on an Intel Core i7-2600 Desktop with 24GB RAM, running Ubuntu 18.04.1 LTS. The scripts and the data used in this analysis are available in the supplementary dataset.

#### 4.1.2 Pathway analysis on S. cerevisiae and P. stipitis joint metabolic network

To identify the metabolic exchanges happening between *S. cerevisiae* and *P. stipitis*, we first enumerated all the pathways till the length 75 using the *find*_*pathways* function in MetQuest package. We constructed the joint bipartite graph of *S. cerevisiae* and *P. stipitis* from their respective genome-scale metabolic models (33, 35) using the *construct*_*graph* module in MetQuest python package. We used the components of YNB medium as seed metabolites along with D-glucose as the carbon source. Using home-grown Python scripts, we analysed every pathway from *S. cerevisiae* that lead to every *scope* metabolite in *P. stipitis* for the presence of metabolites from *S. cerevisiae*. We carried out these simulations on an Intel(R) Xeon(R) CPU E7-4850 v4 @ 2.10GHz workstation with 1TB RAM, running CentOS 7.5.1804. The scripts and the data used in these analyses are available in the supplementary dataset.

### 4.2 Experimental verifications of the predicted metabolic interactions between S. cerevisiae and P. stipitis

#### 4.2.1 Yeast strains used

In this study, we used *S. cerevisiae* (MTCC No. 171) and *P. stipitis* (NCIM No. 3497), procured from Microbial Type Culture Collection, Chandigarh, India and National Collection of Industrial Microorganisms, Pune respectively.

#### 4.2.2 Culture maintenance and inoculum preparation

The yeast strains were independently maintained on YPD agar consisting of 3 g/L Yeast extract, 10 g/L Peptone, 20 g/L Glucose and 1% agar (HiMedia Laboratories Pvt Ltd, Mumbai, India). From the agar plate, one colony was picked and inoculated in YPD medium and grown at 30°C. For the long-term storage, cultures from mid-log phase were collected and maintained as 30% (v/v) glycerol stocks and stored at - 80°C.

Glycerol stocks of the respective yeast strains were revived by streaking onto a YPD Agar plate and incubating at 30°C for 24 h. The primary cultures were initiated as suspension cultures by inoculating single colony on YPD medium, followed by YNB minimal medium without amino acids (Sigma-Aldrich, Germany) supplemented with 10 g/L glucose (Carl Roth GmBH, Germany). The culture was grown in minimal medium till mid-logarithmic phase and used for subsequent experiments. All the experiments were carried out in YNB minimal medium using glucose (Carl Roth GmBH, Germany) as the carbon source.

#### 4.2.3 Growth kinetics of mono- and co-culture

To determine the substrate utilization and product secretion profiles, growth kinetic experiments of both the yeast strains, *S. cerevisiae* and *P. stipitis* were carried out. The culture from the well-grown primary inoculum was inoculated in 500 mL Erlenmeyer flasks consisting of 25 mL of YNB medium with 10 g/L Glucose. For all the experiments, we maintained the starting Optical Density (OD) as 0.17. The experiment was carried out at 300 rpm, 30°C and the growth was continuously monitored online using Cell Growth Quantifier (Aquila Biolabs GmBH). Samples were withdrawn at regular intervals for analyzing the spent metabolites through High-Performance Liquid Chromatography (HPLC).

#### 4.2.4 Growth analysis in the spent medium for checking metabolic interactions

We designed a two-step shake flask experiment to determine if there were any interactions between the organisms. Briefly, in the first stage, *S. cerevisiae* and *P. stipitis* were independently grown in minimal YNB medium with 10 g/L Glucose. The growth was continuously monitored online using Cell Growth Quantifier (CGQ) (Aquila Biolabs GmBH), and the samples were withdrawn at the start of the experiment, early and late exponential phase. CGQ measures the backscattered light emitted by the growing microbial cells. Data analysis, processing and visualisation, were carried out using CGQuant software version 7.3 (Aquila Biolabs GmBH).

Samples were analysed in HPLC to check if the carbon source was completely depleted. At this stage, the cells were separated from the broth by centrifuging at 8000 rpm, 4°C for 10 minutes and the supernatant was filter sterilised. In the next step, to these supernatants, YNB medium and 5 g/L glucose were added. The working volume was 20 mL. We designated the spent medium obtained from *S. cerevisiae* and *P. stipitis* as *SupSce* and *SupPst* respectively. We inoculated these supernatants with either of these organisms, i.e., to the *SupSce* we inoculated *P. stipitis* and to the *SupPst* we inoculated *S. cerevisiae*, such that the initial Optical Density was 0.17 ± 0.1. We continuously monitored the growth of the organisms using CGQ and analysed the samples for residual carbon source and the extracellular metabolites using HPLC at the start and end of experiments. Also, we compared the growth of the organisms with that in the control, where the supernatant was replaced with sterile distilled water. The experimental scheme is as shown in Figure 2.

To check for the metabolic interactions, we repeated the same two-stage experiments as above, with a single modification. Instead of naturally labelled glucose, we used 100% uniformly labelled (U-^13^C) glucose as the carbon source in the first step to cultivate the micro-organisms individually. Samples withdrawn were analyzed through HPLC for the residual carbon and the extracellular metabolites. The biomass at the end of 24 h was analyzed using Gas Chromatography-Mass Spectrometry (GC-MS) to identify the labelling patterns of the amino acids alanine, valine, serine and aspartate.

#### 4.2.5 Analytical methods

##### 4.2.5.1 Quantification of glucose and other by-products

The substrate and the by-product analytes from the sample were estimated using High Performance Liquid Chromatography (HPLC) (System Gold125 Solvent Module, Beckman Coulter, USA) with an Organic acid stationary phase (300 x 8 mm, 10µm particle size) (CS Chromatographie Service GmbH, Langerwehe, Germany) under the following conditions: Mobile phase - 5 mM Sulfuric acid, Flow rate - 0.5 mL/min, Column Temperature - 50 °C using a Refractive Index Detector (Company). Standard plots (with R^2^ = 0.98) for the metabolites ethanol (Carl Roth GmbH, Germany) (0.25 – 2 % (v/v)) and glucose (2-15 g/L) were prepared by injecting known quantities and measuring the area under the respective peaks (Supplementary Figures S3, S4).

##### 4.2.5.2 GC-MS to identify labelling patterns in biomass and identify the metabolites that have been exchanged

To identify the labelling patterns in the biomass, we adapted the protocol from (37). Briefly, 0.3-0.4 mg of the biomass pellets were resuspended in 150 µL 6M Hydrochloric acid (HCl) and transferred to 1.5 mL glass vials (Part No AR0-3940-12 phenomenex). The suspension was incubated at 105°C for 6 hours for hydrolysis, and dried overnight at 85°C. To the dried hydrolysate, 30µL acetonitrile and 30µL *N*-methyl-*N*-tert-butyldimethylsilyl-trifluoroacetamide (MBDSTFA) was added and incubated at 85°C for 1 h. The samples were cooled and immediately analysed using a GC-MS single quadrupole system using the ‘Full scan’ mode. The system consisted of a TRACE™ GC Ultra, an ISQ single quadrupole MS with electron impact ionisation (Thermo Fisher Scientific, Waltham, MA, USA) and a ThermoScientific TriPlus RSH Autosampler. The separation of the amino acids was achieved using TraceGOLD TG-5SilMS fused silica column (length 15 m; inner diameter 0.25 mm; film thickness 0.25 µm). The injector temperature was set at 270°C, and the column oven was set at 140°C for 1 minute, and the temperature was steadily increased to 310°C with a ramp of 10°C/min, with a hold time of 1 minute. The equipment was operated under a steady gas flow of 1 mL/min of Helium with a split ratio of 1/15. For every measurement, 1 µL of the sample was injected. The resulting chromatogram and the mass spectra of the samples were analysed by comparing with those of amino acids standards (Sigma Aldrich, Germany). All the mass spectra and the chromatograms were analysed out on Thermo XCalibur 2.2 software (Thermo Fisher Scientific, Waltham, MA, USA).

##### 4.2.5.3 In silico methods to identify the isotopomer distribution in amino acids

From the chromatogram and the mass spectra, the peaks were identified by comparing the retention times of different amino acids with that of standards. The mass spectrum of each derivatised amino acid was also compared against the NIST Library using NIST MS Search software (https://www.nist.gov/srd/nist-standard-reference-database-1a-v17) to confirm the presence of corresponding amino acid. Also, we determined the average carbon labelling and the relative abundance of every fragment after performing corrections for proton gain and original biomass using iMS2Flux software (Poskar et al., 2012). For all our calculations, we used the (M-57) fragment of the amino acids for analysis.

## ACKNOWLEDGEMENTS

The authors thank Dr Suresh Sudarsan, DTU Biosustain, Novo Nordisk Foundation Center for Biosustainability and Dr Ulf Liebal, iAMB, RWTH Aachen for the useful discussions and critical comments. A.R. acknowledges the funding from the Bundesministerium fur Bildung und Forschung (BMBF), Federal Republic of Germany and the University Grants Commission (UGC), Government of India (A.R.) for funding the exchange program and the INSPIRE fellowship, Department of Science and Technology (DST), Government of India.

## DECLARATIONS OF INTEREST

None

